# Improved detection and phylogenetic analysis of plant proteins containing LysM domains

**DOI:** 10.1101/2023.06.21.545963

**Authors:** Dardo Dallachiesa, O. Mario Aguilar, Mauricio J Lozano

## Abstract

Plants perceive N-acetyl-d-glucosamine-containing oligosaccharides that play a role in the interaction with bacteria and fungi, both pathogenic and symbiotic, through cell-surface receptors that belong to the Receptor-Like Kinase (RLK) or Receptor-Like Protein (RLP) families. Structurally characterised proteins from these families have been shown to contain a tight bundle of three LysM domains in their extracellular domain. However, the identification of LysM domains of RLK/Ps using sequence based methods has led to some ambiguity, as some proteins have been annotated with one or only two LysM domains. This missing annotation was likely produced by the failure of the LysM hidden Markov model (HMM) from the PFAM database to correctly identify some LysM domains in proteins of plant origin. In this work, we provide improved HMMs for LysM domain detection in plants, that were built from the structural alignment of manually curated LysM domain structures from PDB and AlphaFold. Furthermore, we evaluated different sets of ligand-specific HMMs that were able to correctly classify a limited set of fully characterised RLK/Ps by their ligand specificity. In contrast, the phylogenetic analysis of the extracellular region of RLK/Ps, or of their individual LysM domains, was unable to discriminate these proteins by their ligand specificity. The HMMs reported here will allow a more sensitive detection of plant proteins containing LysM domains and help improve their characterisation.

## INTRODUCTION

Plants perceive molecular signals from either pathogenic or mutualistic interactions in which cell-surface receptors, such as LysM containing proteins, play key roles. The lysin motif (LysM) domain is a widespread protein module involved in binding peptidoglycan (PGN), chitin, and lipo-chitooligosaccharides (LCOs), most likely by recognising their common N-acetyl-d-glucosamine (NAG) moiety (Buist et al. 2008). It has a length of 40 to 65 amino acids, and it can be found in different architectures (Bateman and Bycroft 2000). The LysM domain was originally identified in the lysozyme of *Bacillus* phage φ29 (Buist et al. 2008), and since then it has been found in a wide diversity of enzymes, both in prokaryotes and eukaryotes, including PGN hydrolases (Vermassen et al. 2019), peptidases (Razew et al. 2022), esterases, reductases, nucleotidases (Bhaduri and Sowdhamini 2005), and chitinases (Onaga and Taira 2008). LysM domain containing proteins are also involved in bacterial pathogenesis, acting as antigens, and binding to albumin, elastin or immunoglobulins (Bateman and Bycroft 2000; Buist et al. 2008). In eukaryotes, the distribution of the LysM domain is irregular, and it has been mostly found in organisms that tightly interact with bacteria, which suggested a possible horizontal gene transfer origin (Bateman and Bycroft 2000). Unlike microbial LysM domain containing proteins, usually linked to secretory enzymatic activities which modify NAG-containing substrates, most of plant LysM domain proteins characterised to date either have chitinase activity (Ohnuma et al. 2008) or recognise NAG-containing oligosaccharides that play a role in the interaction with bacteria and fungi, both pathogenic and symbiotic (Gust et al. 2012; Broghammer et al. 2012). Plant LysM domains are present in the extracellular Domain (ECD) of proteins belonging to specific Receptor-Like Kinase (RLK) and Receptor-Like Protein (RLP) families (Gough and Cullimore 2011; Gust et al. 2012; Zipfel and Oldroyd 2017). Based on their intracellular kinase domains two main types of plant LysM-RLKs have been defined: LYKs, which have a canonical RD kinase domain and show *in vitro* autophosphorylation activities; and LYRs, which carry an aberrant kinase domain and do not exhibit auto-phosphorylation or trans-phosphorylation activities when tested *in vitro*. In addition, there are GPI-anchored LysM-RLPs that lack the intracellular kinase region and are called LYMs (Buendia et al. 2018). RLKs and RLPs always contain three extracellular LysM domains that are usually quite different in sequence (Arrighi et al. 2006; Bensmihen et al. 2011) and are packed tightly together by disulphide bonds formed by highly conserved cysteine pairs separated by one amino acid (CXC). The formation of these bonds has been shown to be essential for nod factor perception (Miya et al. 2007; Buendia et al. 2018). Plant LysM domains also display extensive, though not completely conserved, N-glycosylation which in certain cases proved not to be required for their biological function (Lefebvre et al. 2012). Finally, RLK/Ps have been shown to form dimers or multimers, involving both LYKs and LYRs, that are required for the signal transduction and regulation (Madsen et al. 2011; Wan et al. 2012; Cao et al. 2014; Han et al. 2014; Fliegmann et al. 2016; Feng et al. 2019).

In legume plants, the nod factors secreted by rhizobia are essential signalling molecules for the establishment of the nitrogen fixing symbiotic interaction that leads to nodule formation. Nodulation factors are sensed by heterodimers of RLKs identified as NFP/Lyk3 and NFR5/NFR1 that contain ECD LysM domains (Moling et al. 2014; Poole et al. 2018; Roy et al. 2020). Nevertheless, the role of the different LysM domains in ligand binding specificity is not completely clear. In several reports, potential CO (Liu et al. 2012; Bozsoki et al. 2017), or LCO binding pockets (Radutoiu et al. 2007; Bensmihen et al. 2011) were identified in the LysM2 domain, which also showed to be important for the discrimination of LCOs from different rhizobia. Furthermore, LysM1, LysM3, and pockets formed by the interaction of LysM domains in RLK/Ps dimers have also been proposed to be important for the ligand recognition and binding. Nod factor and chitin selectivity was reported to be determined by the LysM1 domain (Bozsoki et al. 2020). The LysM3 domain was reported to be important for the high-affinity binding of symbiotic LCOs by most LYR3 proteins, presenting two potential hydrophobic tunnels which were predicted to be able to accommodate the LCO acyl chain (Malkov et al. 2016). LYR3 was also proposed to be involved in the LCO-dependent regulation of MtLYK3 during both nodulation and mycorrhization (Fliegmann et al. 2016). Chitin oligomers were found to induce AtCERK1 ECD dimerization, which is critical for its activity, by acting as a bivalent ligand (Liu et al. 2012). Through the Molecular Modeling of the interaction between RLK heterodimers it was proposed that NFs were recognised by their fatty acid tail in the hydrophobic pocket formed between the LysM domains of the interacting proteins (Igolkina et al. 2018; Solovev et al. 2021). Lastly, in the case of *Lotus japonicus-Mesorhizobium loti* symbiosis, exopolysaccharides sensing has also been demonstrated to occur through the LysM domain containing exopolysaccharide receptor 3 (EPR3) ECD (Wong et al. 2020).

Although the importance of RLK/Ps has been substantially demonstrated for plant immunity and symbiotic interactions, there is some ambiguity regarding the architecture of its domains as determined by *in silico* analysis. Within this context, by browsing the UNIPROT database, we found some RLK/Ps to be annotated with a single or only two LysM domains (https://www.uniprot.org/; e.g. Q70KR8, Q70KR1, Q0GXS4, D3KTZ2; accessed 1/6/2022). We concluded that this missing annotation was produced by the failure of the PFAM database LysM hidden Markov model (HMM) to correctly identify some of the plant LysM domains. This error has unfortunately propagated to other databases, and also generated certain imprecise reports of LysM domain architecture in plant RLKs (Zhang et al. 2007, 2009). Therefore, in this work we built improved HMMs for LysM domain detection based on the structural alignment of manually curated LysM domain structures from PDB and AlphaFold. We also defined HMMs from the structural alignment of triple LysM bundles from plant RLK/Ps, for their LysM1, LysM2, and LysM3 domains, and for their reported ligand binding regions. Using these models we attempted the classification of RLK/Ps by their affinity for PGN, CO and LCOs. In addition, we performed phylogenetic analyses of individual LysM domains that in certain cases show a separation between proteins that are associated with the ligand LCO or CO, respectively. Moreover, we found that, in most cases, the use of ligand specific HMMs was able to correctly classify the RLK/Ps by their ligand specificity. Overall, the HMMs here reported could help improve the detection and characterisation of plant proteins containing LysM domains, although a larger number of RLK/Ps characterised at the ligand level is necessary to improve our findings.

## MATERIAL AND METHODS

### Construction of LysM domain Hidden Markov Models

In order to build new HMMs for the identification of LysM domains, a search for homologous domains with three-dimensional structure was conducted by means of the hmmsearch program from the HMMER3 suit (Eddy 2011) using the PFAM (Finn et al. 2014) LysM domain HMM (PF01476, accessed 14/6/2022) as query, and the amino acid sequences of proteins with 3D structure from the PDB database as target. LysM domain structures were extracted from PDB files using homemade python scripts and the Bio (Cock et al. 2009), Prody (https://github.com/prody/ProDy) and Pandas (https://pypi.org/project/pandas/) modules. Next a multiple structural alignment was done with mTm-align (Fig. 1S) (Dong et al. 2018), and the corresponding sequence alignment was used to build a HMM with the hmmbuild tool (HMMER3, LysM-PDB HMM). This model was used to search for LysM domain containing proteins, obtaining results that were comparable –in numbers– to those previously obtained with the PFAM model. Since our HMM could be further improved, a LysM blast search was conducted to look for LysM domains in the AlphaFold database (Varadi et al. 2022) (https://alphafold.ebi.ac.uk/), and the structural models of those homologs were downloaded and added to the structural alignment. This alignment was manually curated to eliminate misaligned domains, and a total of 273 aligned structures were used to build a new HMM (Fig. 1.A; Table 1S). Since some of the proteins identified by the PFAM HMM were missing (Reference proteomes –PFAM, PF01476– Accessed 13-06-2022), they were added to the alignment using the MAFFT aligner with the --add option (MAFFTv7, Katoh and Standley, 2013) a new HMM –LysM-ST– was built (Fig. 1.B, Table 1S, https://github.com/maurijlozano/Plant-LysM-Domain-HMM,LysM-ST.hmm).

**Figure 1.**
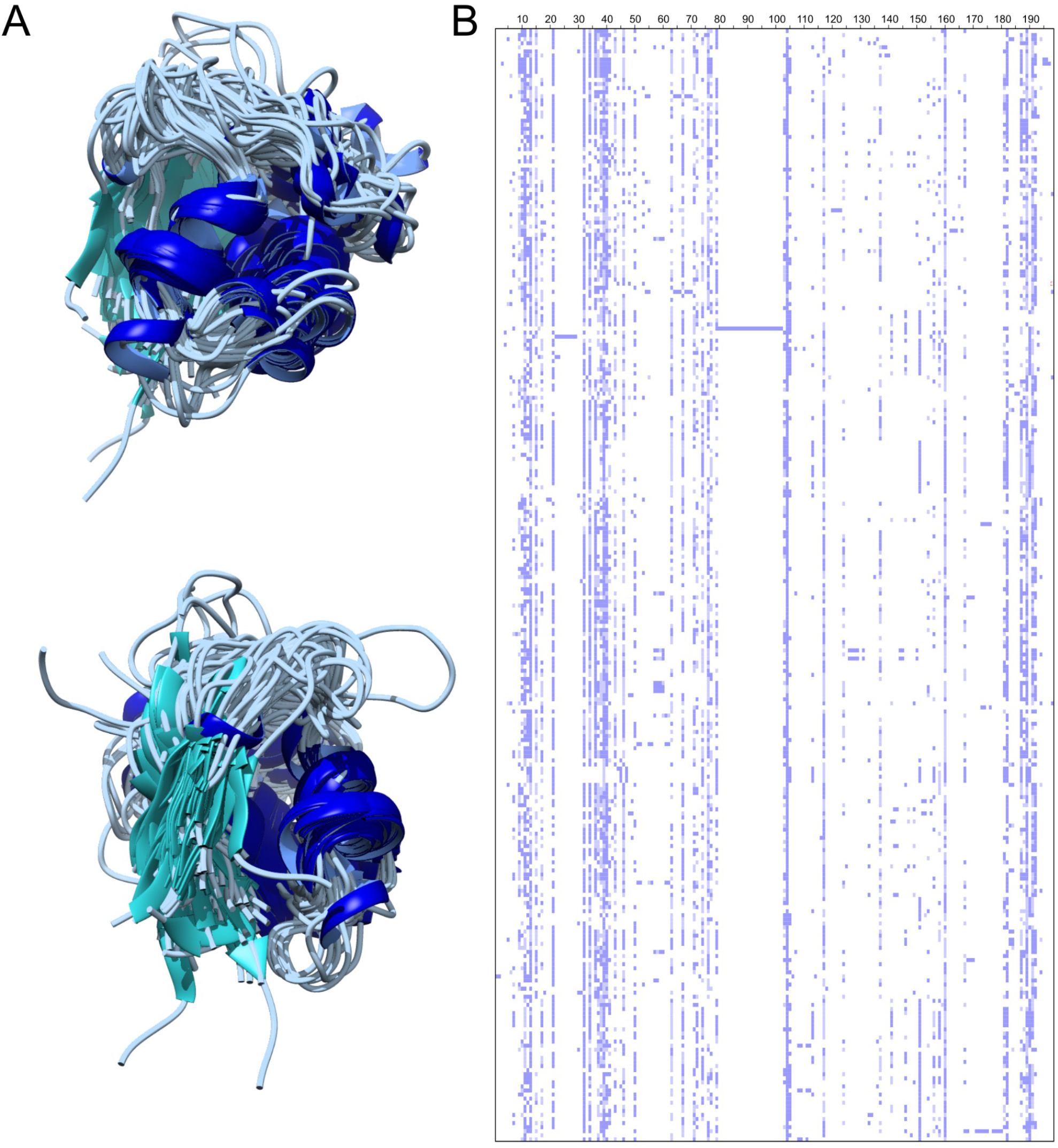
Structural alignment of LysM domains. A. Superposition of the 273 structures from PDB and AlphaFoldDB that were used in the structural alignment. B. Schematic view of the sequence alignment derived from the LysM structural alignment that was used to build the LysM-ST HMM. The list of sequences used in the alignment can be found in Table 1S.

In addition, a HMM of the triple LysM domain bundle present in plant RLK/Ps was constructed. To that end, a first structural alignment of 5LS2, 6XWE, 7AU7, 7BAX crystallographic structures was made using mTm-align and the derived multiple sequence alignment was used to build a HMM. This model was used to look for proteins in the PDB database. In addition, 7BAX structure was used to look for homologs in the AlphaFold database using Foldseek (van Kempen et al. 2023) server (https://search.foldseek.com/search). A multiple structural alignment was constructed with a total 48 structures and AlphaFold models using mTm-align (Fig. 2.A). Six misaligned structures were removed, and 32 additional sequences were added to the derived sequence alignment (Fig. 2.B) using mafft with the --add option, and the resulting alignment was used to build a HMM (LysM-3B, https://github.com/maurijlozano/Plant-LysM-Domain-HMM/tree/main, LysM-3B.hmm). In addition, the LysM1, LysM2 and LysM3 domains were extracted from the previous alignment and used to build the corresponding HMMs.

**Figure 2.**
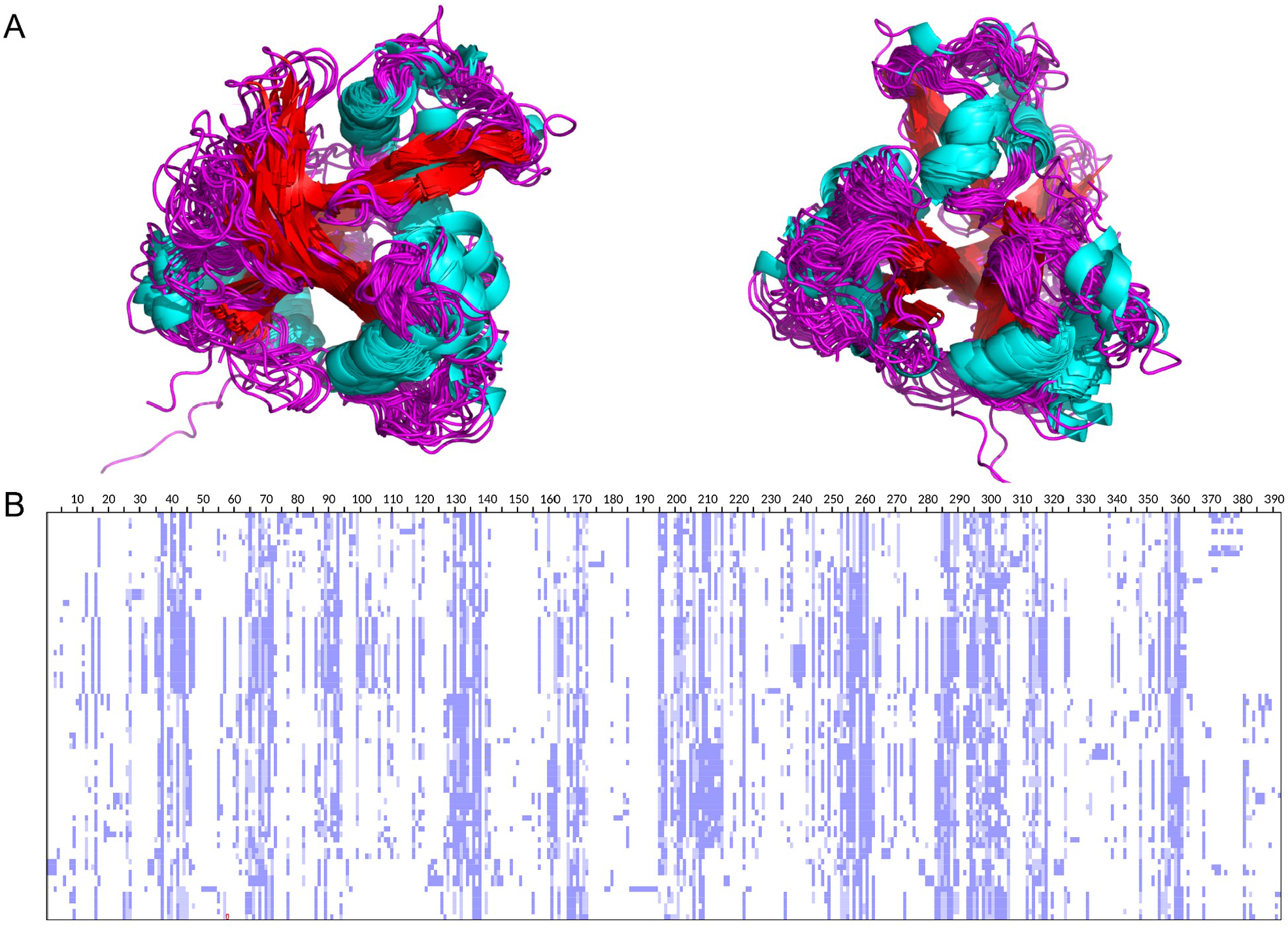
Structural alignment of plant LysM triple bundle domains. A. Superposition of the 48 structures from PDB and AlphaFoldDB used in the structural alignment. B. Schematic view of the sequence alignment derived from the LysM triple bundle domain structural alignment that was used to build the LysM-3B HMM. The list of sequences used in the alignment can be found in Table 1S.

The sequences with experimentally determined ligand specificity were extracted from the LysM-3B alignment and classified by their ligand (Fig. 3). Three sets of ligand specific HMMs were built, one for the triple LysM domain bundle, one for the individual LysM domains, and one for the previously reported NAG-oligosaccharides (CO/LCO/PGN) binding regions (Radutoiu et al. 2007; Ohnuma et al. 2008; Bensmihen et al. 2011; Liu et al. 2012; Mesnage et al. 2014; Malkov et al. 2016; Bozsoki et al. 2017, 2020; Solovev et al. 2021). In addition to the residues reported to be in contact with the ligand, the loops between *β*_1_-α_1_ and α_2_-*β*_2_ were taken into account. Sequence alignments were visualised and cured in Jalview v2 (Waterhouse et al. 2009). To evaluate the performance of the HMMs the EBI-HMMER server was used (https://www.ebi.ac.uk/Tools/hmmer/). Tridimensional models and multiple structural alignments were rendered with Pymol (Schrödinger 2010) or UCSF Chimera (Pettersen et al. 2004). When required, structural models were obtained with ColabFold (Mirdita et al. 2022).

**Figure 3.**
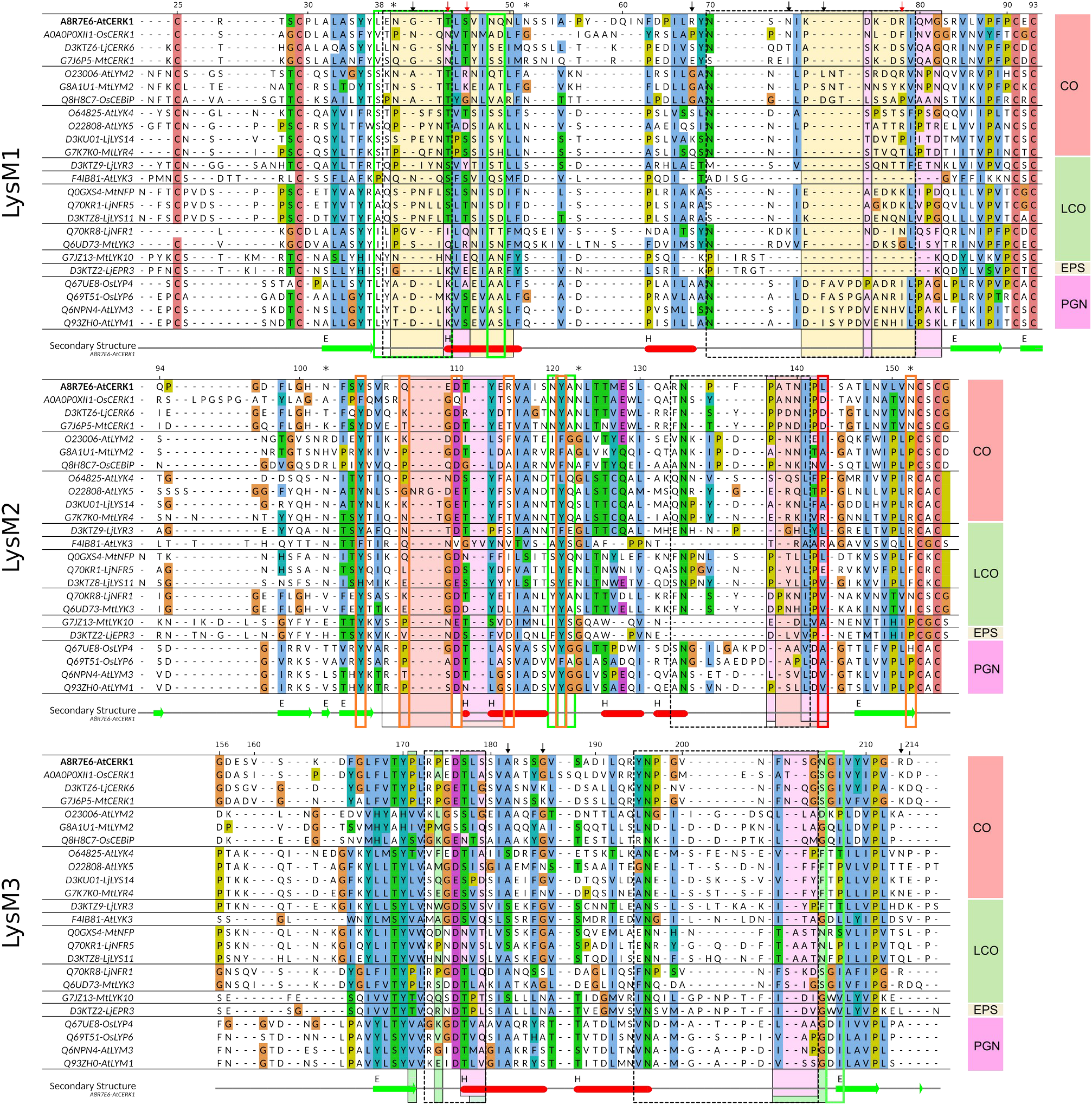
Multiple sequence alignment showing the CO/LCO/PGN binding regions reported in previous works. A selection of 24 sequences from the dataset used by Buendía et al. (2018) was used. Orange shading: regions II and IV from Bozsoki et al. (2020). Red shading, regions involved in CO sensing reported by Bozsoki et al. (2017) and Liu et al. (2012). Magenta: PGN binding region defined in Mesnage et al. (2014). Thick lined red boxes, positions involved in NF discrimination in Bensmihen et al (2011). Thick lined orange boxes, positions reported by Radutoiu et al. (2007). Dashed thick lined black boxes: CO binding site reported by Ohnuma et al. (2008). Thick lined green boxes: positions involved in NF binding reported by Solovev et al. (2021). Green shading: positions involved in NF binding reported by Malkov et al. (2016). Reported * Glycosylation sites in AtCERK1. Arrows: Positions under positive selection (Hyphy MEME, this work). Red arrows: positions under positive selection (De Mita et al. 2014)

### Molecular phylogenetic analysis

Protein sequences from the sequence data-set used by Buendia et al. (2018) were aligned to the structurally based sequence alignment of the triple LysM bundles using the Mafft --add mode, and then the original sequences used to build the HMM were removed (Fig. 2S). The LysM-3B, LysM1, LysM2 and LysM3 sub-alignments were extracted and used to build Phylogenetic trees using IQ-Tree (Minh et al. 2013; Nguyen et al. 2015), with automatic detection of the best evolutionary model, and 1000 ultrafast bootstrap replicants. In addition, proteins with known ligand binding specificity were selected from the Buendia et al. (2018) dataset, their structural models recovered from AlphaFoldDB and a multiple structural alignment of the ECDs was obtained with mTm-align. LysM domains were identified, and their corresponding sequence alignments were used to build Phylogenetic trees using IQ-Tree with automatic detection of the best evolutionary model, and 1000 ultrafast bootstrap replicants. All the tree figures were generated using either ITOL (Letunic and Bork 2019) or Figtree (Rambaut and Drummond 2009), and edited with Inkscape (TEAM-Inkscape) when required. Positive selection was determined using the program MEME from the Hyphy suite (Suyama et al. 2006) at the Datamonkey adaptive evolution server (https://www.datamonkey.org/meme). For this analysis, a codon alignment was generated from the amino acid sequence alignment and the corresponding coding sequences using Pal2nal (Suyama et al. 2006). Interprotein coevolution signals were analysed with I-COMS (Iserte et al., 2015) using the plmDCA score. A circos plot with the 150 top links is shown in figure 4S.B.

## RESULTS

### Revisited LysM HMM profiles improve the identification of plant LysM domains

The PFAM LysM (PF01476) HMM failed to detect a wide range of LysM domains in plant RLK/Ps, missing in general the detection of LysM1 (and in some cases LysM2, Table 2S) domains. To improve LysM detection in plants we defined a set of new HMMs based on the structural alignment of LysM domains with known tridimensional structure, including plant LysM domain containing proteins. First, we built a new general HMM (LysM-ST) from the structural alignment of LysM domains from X-ray crystallography and AlphaFold structural models. To evaluate the sensitivity of the new model, we used hmmsearch locally to look for both, the PFAM LysM HMM (PF01476, LysM-PFAM) and the LysM-ST HMM using UNIREF50 as protein database (Table 1). In general, the performance of the LysM-ST HMM resulted comparable to that of the LysM-PFAM HMM (c.a. the same number of recovered homologs) but some differences were found, with the LysM-ST performing better for plant sequences (Table 1), being able to successfully recover the three LysM domains in most of the RLK/Ps (Table 1, 65 % / 92 %).

**Table 1.** HMMSearch results.

| DATASET |  | HMM |  |  |  |
| --- | --- | --- | --- | --- | --- |
| UNIREF50 Release 2023_01 (59,142,917 Protein representatives) | hmmsearch options | LysM-PFAM | LysM-ST | LysM-3B |  |
| Domains (Filtered by I-Evalue < 0.001) | --domE 0.001 | 47290 | **47774** | - |  |
| Domains (Filtered by I-Evalue < 0.001, with "LysM" in Protein description) |  | 34176 | **34747** | - |  |
| Domains (Filtered by I-Evalue < 0.001, with "LysM" in Protein description) |  | 701 | **980** | - |  |
| Proteins (Filtered by E-value < 0.001) |  | 38089 | **38602** | 13158 |  |
| Proteins (Filtered by E-value < 0.001 and with "LysM" in Protein description) |  | 27759 | **27967** | 9151 |  |
| Proteins (Filtered by E-value < 0.001 and for "*Viridiplantae*") |  | 853 | **1211** | 926 |  |
| UNIPROT-Refprot (v.2021_04) at www.ebi.ac.uk/Tools/hmmer |  |  |  |  |  |
| Proteins (Filtered for "*Viridiplantae*") | default options | 2978 | **4427** | 3234 |  |
| RLK/Ps (123 sequences; Buendia et al. 2018) (Total confirmed LysM: 367)* |  | Total Count | % Recovery | F+ | F- |
| LysM-PFAM ALL Proteins (E-value < 0.001) | --domE 0.001 | 117 | 95,12 | 0 | 6 |
| LysM-PFAM ALL Domains (I-Evalue < 0.001) | --domE 0.001 | 143 | 38,96 | 0 | 224 |
| LysM-PFAM ALL Domains | - | 278 | 75,20 | 2 | 91 |
| LysM-ST ALL Proteins (E-value < 0.001) | --domE 0.001 | **123** | **100,00** | 0 | 0 |
| LysM-ST ALL Domains (I-Evalue < 0.001) | --domE 0.001 | **240** | **65,40** | 0 | 127 |
| LysM-ST ALL Domains | - | **341** | **92,37** | 2 | 30 |
| LysM-3B ALL Proteins (E-value < 0.001) | --domE 0.001 | **123** | **100,00** | 0 | 0 |
| LysM-3B ALL Domains (I-Evalue < 0.001) | --domE 0.001 | **123** | **100,00** | 0 | 0 |
| LysM-3B ALL Domains | - | **126** | **100,00** | 3 | 0 |
| *Confirmed by manual structural inspection at AlphaFoldDB |  |  |  |  |  |
| 3234 sequences with LysM-3B domain, UNIPROT-Refprot Filtered for "*Viridiplantae*" |  | Total Count | % LysM1-3 | % LysM-3B |  |
| Proteins (Detected either by LysM1, LysM2 or LysM3 HMMs) | - | 3188 | - | 98,58 |  |
| Proteins with 3 LysM Domains | - | 2801 | 87,86 | 86,61 |  |
| Proteins with 6 (2*3) LysM Domains | - | 29 | 0,91 | 0,90 |  |
| Correct LysM model order LysM1-LysM2-LysM3 | - | 2778 | 99,18 | 85,90 |  |
| 123 sequences with LysM-3B domain, RLK/Ps, Buendia et al. 2018 |  | Total Count | % LysM1-3 | % LysM-3B |  |
| Proteins (Detected either by LysM1, LysM2 or LysM3 HMMs) | - | 123 |  | 100,00 |  |
| Proteins with 3 LysM Domains | - | 117 | 95,12 | 95,12 |  |
| Correct LysM model order LysM1-LysM2-LysM3 | - | 117 | 100,00 | 95,12 |  |

F+: False positives; F-: False negatives

Next, we built a HMM for the triple LysM bundle (LysM-3B) present in plant RLK/Ps as described in the Materials and Methods section. The results of using this HMM to search the UNIREF50 protein database are shown in Table 1. This HMM was also used to search for RLK/Ps in the hmmsearch web server at the EBI-UK portal (https://www.ebi.ac.uk/Tools/hmmer/) using as database the reference proteomes filtered for plant sequences. In the last case, a total of 3,234 sequences containing the LysM-3B domain were found, which is a significant improvement in comparison with 2,978 proteins with at least one domain found with the PFAM HMM. In addition, LysM1, LysM2 and LysM3 regions were obtained from the LysM-3B structural alignment and used to build the respective HMMs. These HMMs were successfully used to identify LysM1, Lysm2 and LysM3 domains and their relative order in the previously identified plant RLK/Ps (Table 1). 2,801 out of the 3,234 proteins detected by LysM-3B, were identified as having 3 LysM domains, 99.18% of which displayed the correct order LysM1-LysM2-LysM3. For the Buendia et. al. (2018) dataset, 95% of the proteins were identified by the LysM1-3 HMMs as having the 3 LysM domains in the correct order.

Finally, to discriminate on a sequence basis the ligand specificity of RLK/Ps for LCO, CO or PGN, a selection of proteins with experimentally determined ligands (Buendia et al. 2018) were extracted from the LysM-3B structural alignment for further analysis (Table 1S). We used the aligned sequences of the ECD, the three LysM domains, and the regions containing residues known to interact with the ligand in these proteins (Radutoiu et al. 2007; Ohnuma et al. 2008; Bensmihen et al. 2011; Liu et al. 2012; Mesnage et al. 2014; Malkov et al. 2016; Bozsoki et al. 2017, 2020; Solovev et al. 2021)(Fig. 3) to build three sets of HMMs comprising all the combinations of ligand specificities (CO/LCO/PGN) and sequence regions. Using the HMMs for the ECD, the ligand affinity of the 24 RLK/Ps used here was correctly assigned—with the lowest E-value—by the HMM of the corresponding ligand (Table 3S). In the case of the LysM 1, 2 and 3 HMM sets, the three best hits always identified the ligand binding specificity correctly, except in the case of Q6UD73 (MtLYK3). The best hit for CO binding proteins always corresponded to the LysM2-CO HMM. For LCO, the best hit corresponded either to the LysM3-LCO or to the LysM2-LCO HMMs. Lastly, in the case of PGN, it corresponded to the LysM1-PGN or Lys3-PGN HMMs (Table 3S). In all cases Q6UD73 (MtLYK3) was correctly classified by the best hit, but not by the second one. Finally, using the HMMs generated with the religion II and IV of LysM 1, 2 and 3 of proteins with CO, LCO or PGN ligand specificity (Fig. 3), the sequences were correctly classified by at least 2 of the HMMs. Remarkably, in most cases, the best matching HMM corresponded to the correct ligand (Table 3S) with most of the incorrect hits corresponding to the region II from LysM2 and LysM3 CO-LCO HMMs. Using only LysM1 motifs, a total of 21 sequences were correctly classified by their ligand specificity. With the exception of OsCEBiP and AtLYK5, which were correctly classified by the region IV HMMs (region II HMMs couldn’t be found), the remaining sequences were successfully classified by the region II LCO/CO/PGN HMMs.

### Maximum Likelihood phylogenetic analysis was unable to discriminate RLK/Ps by their ligand specificity

A maximum likelihood phylogenetic analysis of plant LysM domains was carried out to assess their association with CO, LCO or PGN ligand specificity. To that end we used a curated RLK/Ps sequence set obtained from Buendia et al. (2018), which includes RLK/Ps sequences from legume and non-legume plants. These sequences were added to the previously constructed LysM-3B structural alignment with MAFFT --add option, and the respective original sequences were removed. Four multiple sequence alignments were generated as follows, one for the LysM-3B, and one for each of the LysM1, LysM2 and LysM3 domains, and used to infer their phylogenetic relationships. The phylogenetic analysis revealed that the LysM-3B domains (Fig. 4, WAG+G4 evolutionary model) clustered mainly by RLK/P family (LYR, LYK and LYM). Furthermore, the individual phylogenetic analysis of LysM1, LysM2 and LysM3 domains belonging to LYK, LYM and LYR RLK/Ps revealed a pattern separating CO and LCO sensing proteins only in the case of LYR (Fig. 5 and Fig. 3S). In the case of LYK RLKs, this separation couldn’t be observed (Fig. 5 and Fig. 3S). Remarkably, all the LysM domains of LYM RLPs clustered by CO or PGN binding specificity (Fig. 3S). In addition, phylogenetic trees of RLK/Ps with known ligand specificity were constructed for LysM1 and LysM2 domains. In the case of LysM2 domains, it could be observed that they clustered mainly by protein class (LYK-I, LYK-II, etc; Fig. 6.) and not by the type of ligand, while for LysM1 most of the domains related to CO sensing were in one clade.

**Figure 4.**
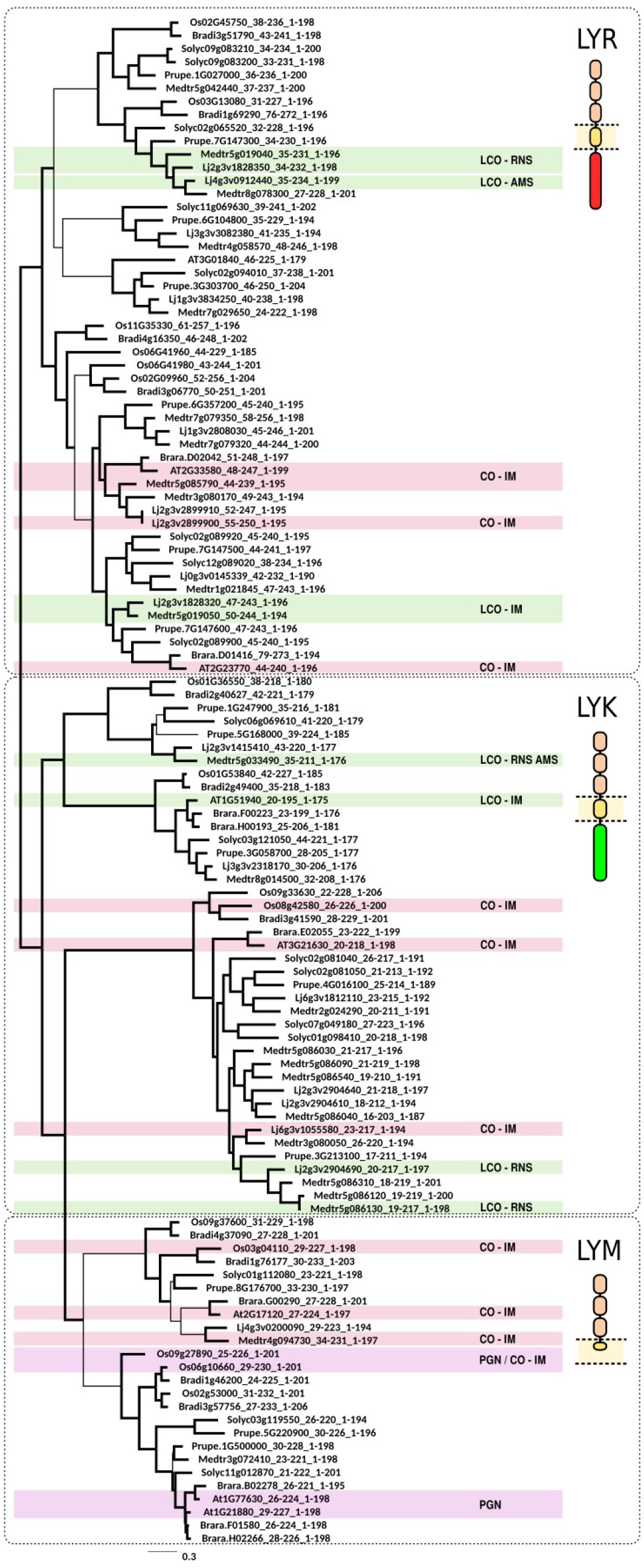
Phylogenetic tree of the extracellular domain of RLK/Ps. A maximum likelihood phylogenetic tree constructed using the sequences of the LysM triple bundles from the extracellular domain of the RLK/Ps used by Buendía et al (2018). CO-IM: LysM triple bundles from Proteins involved in plant immunity triggered by chitin oligosaccharides. LCO-RNS: LysM triple bundles from proteins involved in rhizobium NF perception. PGN: LysM triple bundles from proteins involved in peptidoglycan sensing. Thick lines: Bootstrap support greater than 70%.

**Figure 5.**
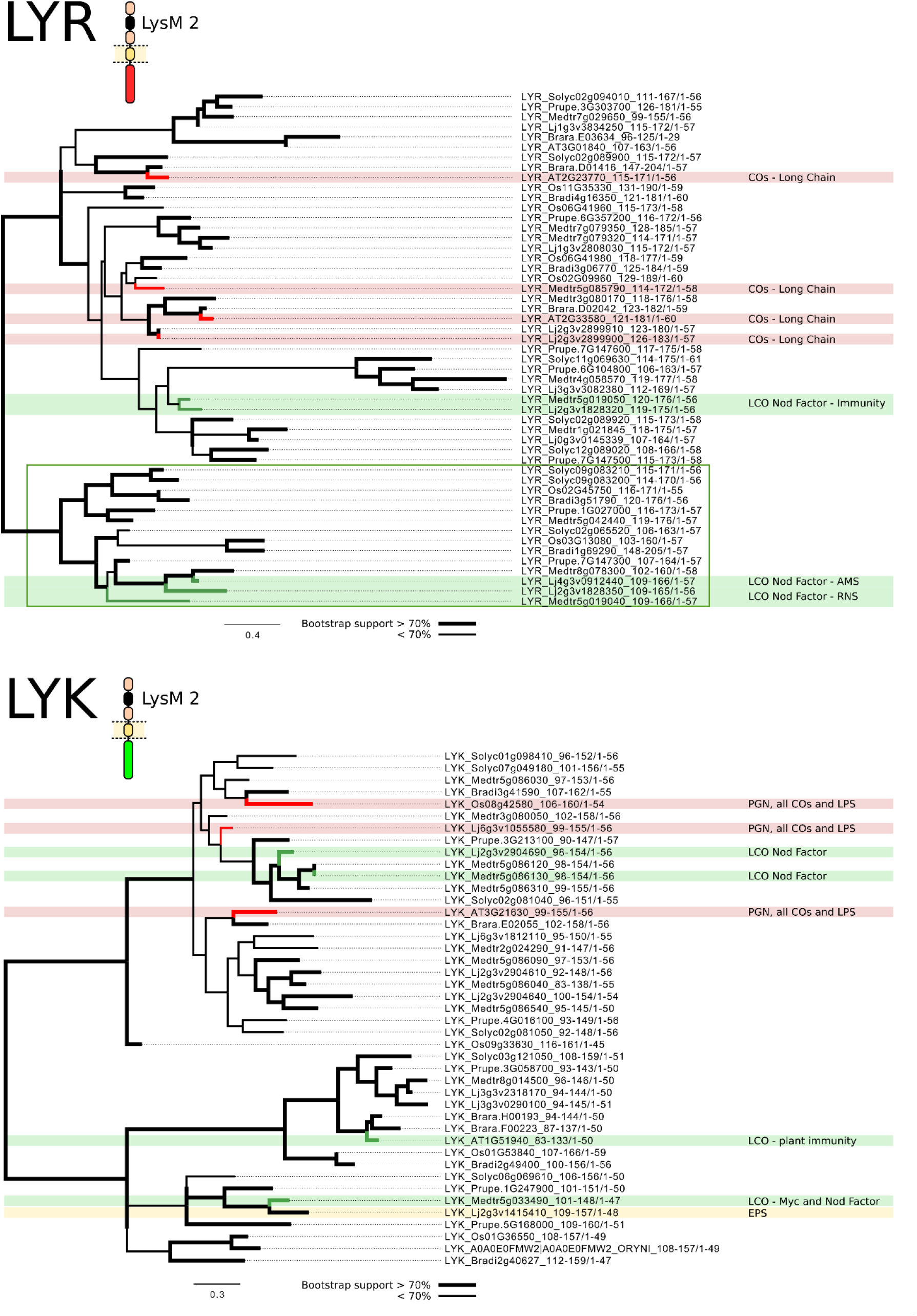
Phylogenetic tree of individual LysM2 domains from LIR and LYK RLKs. Maximum likelihood phylogenetic trees constructed using the LysM2 domains from the sequences used by Buendía et al. (2018) classified by family (LYR,LYK,LYM). COs: chitin oligosaccharides. LPS: lipopolysaccharides. LCOs: Lipo-chito-oligosaccharides from rhizobium (RNS) and mycorrhiza (AMS). PGN: peptidoglycan. EPS: exopolysaccharides. Thick lines: Bootstrap support greater than 70%.

**Figure 6.**
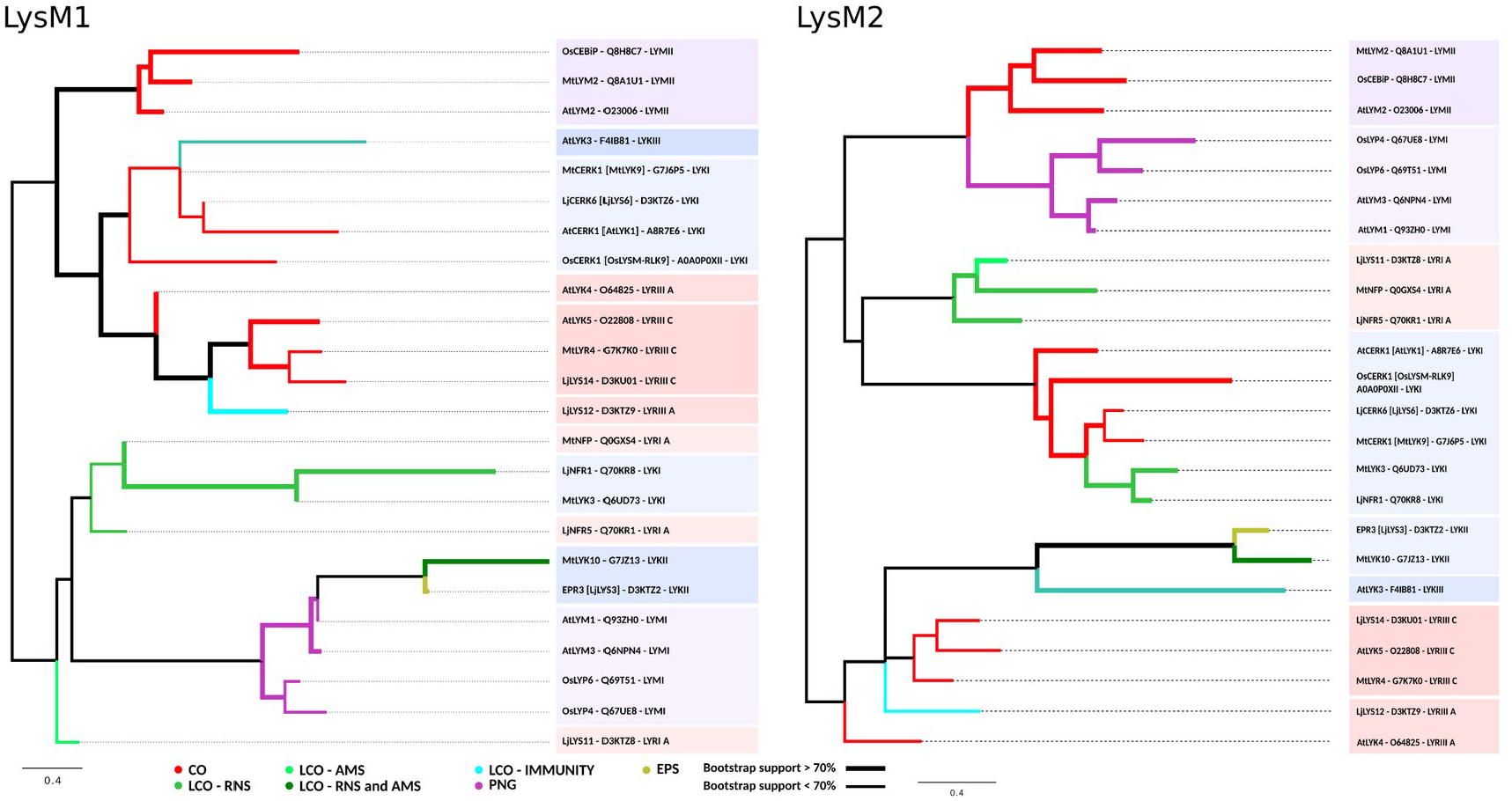
Phylogenetic tree of individual LysM1 and LysM2 domains of RLK/Ps characterised at the ligand level. Maximum likelihood phylogenetic trees constructed using either LysM1 or LysM2 domains from the sequences of RLK/Ps used by Buendía et al. (2018), and with a known ligand specificity. COs: chitin oligosaccharides. LPS: lipopolysaccharides. LCOs: Lipo-chito-oligosaccharides from rhizobium (RNS) and mycorrhiza (AMS). PGN: peptidoglycan. EPS: exopolysaccharides. Thick lines: Bootstrap support greater than 70%.

## DISCUSSION

LysM domains are NAG-oligosaccharide binding domains, that show diverse ligand specificity, being able to bind chitin oligosaccharides, peptide-glycan, and lipo-chito-oligosaccharides, including the Nod factors (NF) secreted by rhizobia. NFs are key molecules involved in the early molecular dialogue between legumes and rhizobia, triggering the formation of N_2_ fixing root nodules (Jones et al. 2007; Oldroyd 2013; Poole et al. 2018; Roy et al. 2020). While studying legume plant RLK/Ps involved in symbiosis, we noticed that the PFAM LysM HMM failed to detect a wide range of LysM domains, missing in most cases the detection of LysM1 (Table 1S). To improve detection of LysM domains of plants we defined a new set of HMMs (LsyM-ST and LysM-3B), which enabled us to detect a significantly higher number of LysM containing proteins (Table 1). Legume plants often have genomic clusters with numerous RLK/Ps (Arrighi et al. 2006; Lohmann et al. 2010; Buendia et al. 2018) making difficult to determine the true orthologs. Since our primary interest was to detect RLK/Ps that might act as NF receptors, we looked for sequence signatures that might be related to ligand binding specificity. One of the limitations that we faced was that there was a low number of RLK/Ps characterised at the ligand level. We attempted a phylogenetic classification using the RLK/Ps ECDs. However, it could be seen that the sequences clustered mainly by RLK/P family (LYR, LYK and LYM, Fig. 4), suggesting, as it was previously reported (Arrighi et al. 2006; Gough et al. 2018), that these sequences must have first diversified from an ancestor RLK/P into different families with CO binding capability (Nakagawa et al. 2011), and then specialised to sense the different ligands. That the common ancestor could have been a CO binding RLK/P is still not completely clear as phylogenetic evidence point that it could have been involved in symbiotic perception of endomycorrhizae (De Mita et al. 2014). In addition, each of the LysM domains were individually analysed, separating them by RLK/P class (LYK, LYM and LYR). This analysis revealed a clear pattern which grouped LYR proteins that sense CO and LCO (Fig. 5. and Fig. 3S). Although the LysM2 domain has been demonstrated to bind CO, and a conserved ligand binding region was defined in the case of AtCERK1, MtCERK1 (MtLYK9) and LjCERK6 (LjLYS6) (Bozsoki et al. 2017), our analysis showed no clustering by ligand specificity in LYK RLKs. Two diverging motifs in the LysM1 domain, regions II (GSNLTY) and IV (KDSVQA), have been described to be necessary for discriminating between immunity and symbiotic functions (Bozsoki et al. 2020). Although these residues are highly conserved among legume CERKs, we couldn’t see a clear clustering by the type of ligand in the LysM1 domain trees (Fig. 3S). Remarkably, all the LysM domains of LYM-RLPs clustered by CO or PGN binding specificity (Fig. 3S). In particular, the LysM1 domain of PGN sensing RLK/Ps presented a conserved Y[AT]D[LM]K motif in the region II that is absent in the CO/LCO sensing LysM domains (Fig. 3). However, this motif wasn’t conserved in bacterial PGN sensing LysM domains (Mesnage et al. 2014), which could indicate that this feature could be only a consequence of the divergence between RLKs and RLPs.

In addition, we examined the LysM1 and LysM2 domains of RLK/Ps with a known ligand. LysM2 domains clustered mainly by protein class (LYK-I, LYK-II, etc; Fig. 6) and not by the type of ligand. Unlike LysM2, LysM1 grouped sequences related to CO sensing all in one clade, including both LYMs and LYKs (Fig. 6). These results are in agreement with the observation that receptors recognising Nod factors and chitin are more divergent in the LysM1 domain (Bozsoki et al. 2020), and have been under the effect of positive selection (De Mita et al., 2014, this work)(Fig. 3).

Previous reports indicate that RLK/Ps biological function requires the formation of dimers (Liu et al. 2012; Moling et al. 2014). Using (NAG)_5_ as ligand it was shown that AtCERK1 ECD interacts with only three chitin residues. However, higher biological activity was displayed by chitin octamers, which act as bivalent ligands stabilising AtCERK1-ECD dimers (Liu et al. 2012). Igolkina et al. (2018) proposed a “sandwich-like” configuration of the homo/heterodimer subunits in which an inter-chain binding pocket is formed by the LysM1 domain from one receptor, and the LysM2 and LysM3 domains from the other. A recent work by Solovev et al. (2021) suggested that in the case of nod-factors (NF) the signalling specificity of RLK/Ps could be also determined by dimer formation. These authors hypothesise that the role of NFs is to specifically stabilise receptor dimers, providing time for interaction of kinase domains to run the downstream phosphorylation cascade. In addition, Feng et al. have shown that COs from CO4 to CO8 activate both symbiosis and immunity signaling and suggested that a combination of COs and LCOs promotes a symbiotic outcome, while perception of COs alone induces an immune response (Feng et al. 2019). These observations, together with the low number of amino acids involved in ligand binding (Ohnuma et al. 2008; Bensmihen et al. 2011; Liu et al. 2012; Mesnage et al. 2014; Malkov et al. 2016; Bozsoki et al. 2017, 2020; Solovev et al. 2021), could explain why we couldn’t find a strong phylogenetic signal able to discriminate RLK/Ps by their ligand specificity. To add further support to this assumption we modelled the interaction between *Lotus japonicus* NFR1 and NFR5 proteins using ColabFold implementation of AlphaFold in multimer mode and using paired alignments (Mirdita et al. 2022). The best ranked model (Fig. 4S.A) shows that NFR1 LysM1 and LysM2 domains interact with LjNFR5 LysM2 and LysM3, and that the ligand binding regions could be located in, or near, the interacting surface. This is also in accordance with previous reports showing that LysM1 was necessary for ligand discrimination in LjNFR1 (Bozsoki et al. 2020), while in the case of MtNFP/LjNFR5 the LysM2 domain was required (Radutoiu et al. 2007; Bensmihen et al. 2011). Nevertheless, the other four models generated by AlphaFold presented a different form of interaction, on which LysM2 and LysM3 from NFR1 interacted with the LysM2 and LysM3 from NFR5. Moreover, the predicted models for the interaction between NFR1 and NFR5 homologs in *Medicago truncatula, Phaseolus vulgaris* and *Glycine max* also presented substantially different modes of interaction (Fig. 4S.C), and relatively low pLDDT score in the interphase. In addition, we analysed NFR1-NFR5 interprotein coevolution signals with I-COMS (Iserte et al. 2015) (Fig. 4S.B) using the plmDCA score. The obtained results further support a possible interaction between NFR1-LysM1 and NFR5-LysM3, but also interactions with NFR5-LysM1 which were absent in the AlphaFold model.

Although the phylogenetic approach was inconclusive, we attempted at classifying these proteins by their ligand specificity using HMMs that were defined to match the ECD; the LysM1, 2 or 3 domains; or the regions corresponding to the LysM CO/LCO/PGN binding region (Fig. 3) located in the loops between *β*1-α1 and α2-*β*2. We found of interest that, in most cases the HMMs correctly classified the 24 RLK/Ps with known ligand specificity. The LysM HMM that worked the best in the case CO sensing RLK/Ps was the corresponding to the LysM2 domain, which has been usually regarded as the ligand binding domain. It was not the case in LCO and PGN sensing proteins. Moreover, when we used the HMMs of the binding regions, the best were the ones corresponding to LysM1 regions II and IV. LysM1 was also reported to be important for ligand binding, and possibly for ligand specificity, by forming part of the ligand binding site located in the homo/hetero-dimer interface. Although these are encouraging results, due to the limited number of available RLK/Ps with determined ligand specificity, and to their evolutionary relatedness, we are cautious on the use of these HMMs for ligand specificity inference.

## CONCLUSION

RLK/Ps are versatile plant oligosaccharide sensors with important roles in immunity and symbiosis. Here we provide improved HMMs for identification of LysM domain containing proteins in plants, and for their classification by ligand binding specificity. We also show that there is only a weak phylogenetic signal that could be used to discriminate the RLK/Ps by their ligands, which could be found for LysM1, and for LysM2 in the case of LYRs. These results are in accordance with the ability to bind CO that was reported in both symbiotic and immunity associated RLK/Ps, and could indicate that the mechanism for the discriminative recognition of NAG containing ligands in plants most likely seems to occur through hybrid surfaces involving at least two receptors, together with a distinctive temporospatial expression pattern (Moling et al. 2014; Zipfel and Oldroyd 2017).

## Data availability

The HMMs built in this work can be found in Github (https://github.com/maurijlozano/Plant-LysM-Domain-HMM). Further detailed data that support the findings of this study are available from the corresponding author upon request.

## Author contributions statement

M.J.L and D.D performed experiments. M.J.L, and O.M.A and wrote the paper. M.J.L, D.D and O.M.A discussed the design of experiments and revised the manuscript. All authors read and approved the manuscript.

## Acknowledgements

This work has been possible thanks to the support of the National Science and Technology Research Council (Consejo Nacional de Investigaciones Científicas y Técnicas - CONICET, Argentina), the European Union’s Horizon 2020 research and innovation programme (Intelligent Collections of Food Legumes Genetic Resources for European Agrofood Systems, INCREASE, No 862862) and the National University of La Plata (Universidad Nacional de La Plata, Argentina).

