## Supplementary figures for "Improved detection and phylogenetic analysis of plant proteins containing LysM domains"

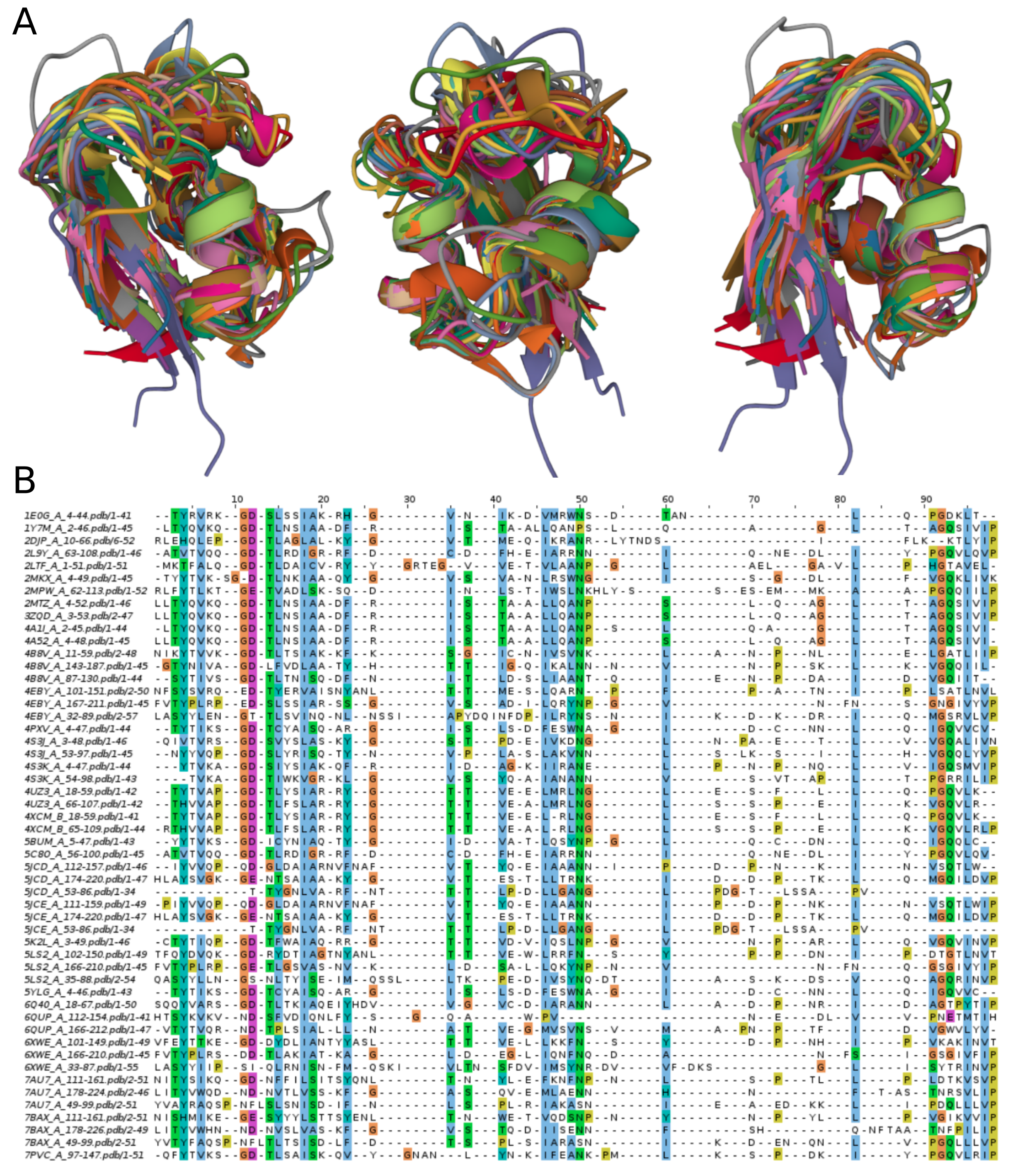
**Figure 1S. Structural alignment of LysM domains from PDB.** A. LysM domain superposition. B. Sequence alignment derived from the structural alignment.

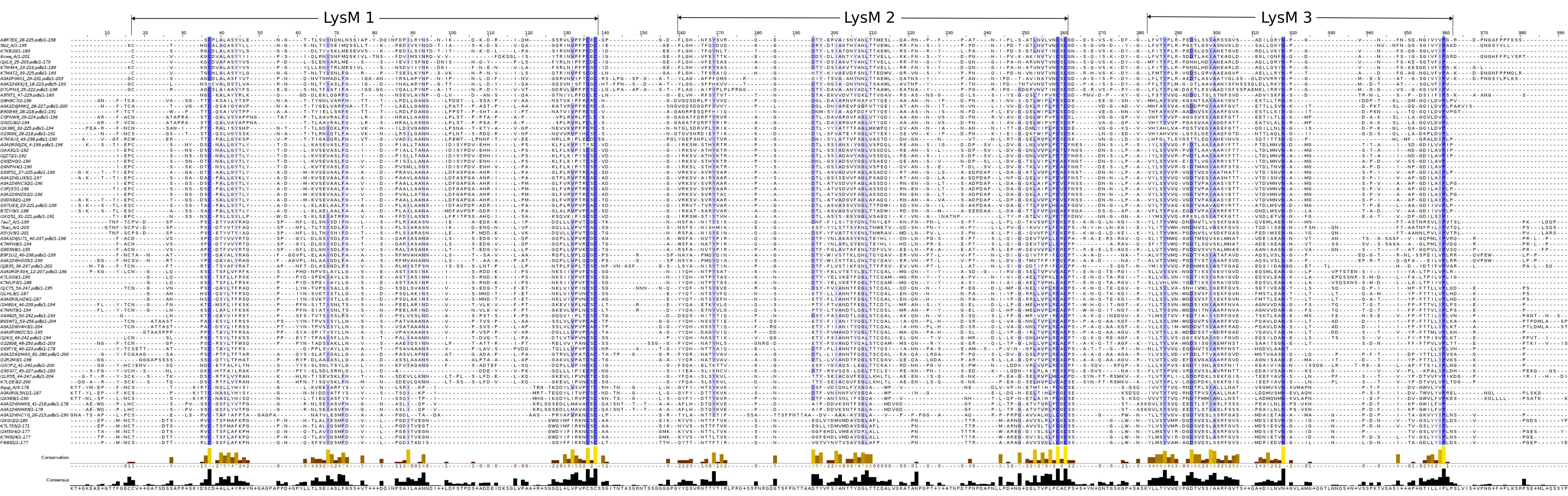
**Figure 2S. Structural alignment of the extracellular domain of RLK/Ps.** Sequence alignment derived from the structural alignment of the extracellular domain of RLK/Ps obtained from the sequences used by Buendía et al. (2018). Blue shading: conservation.

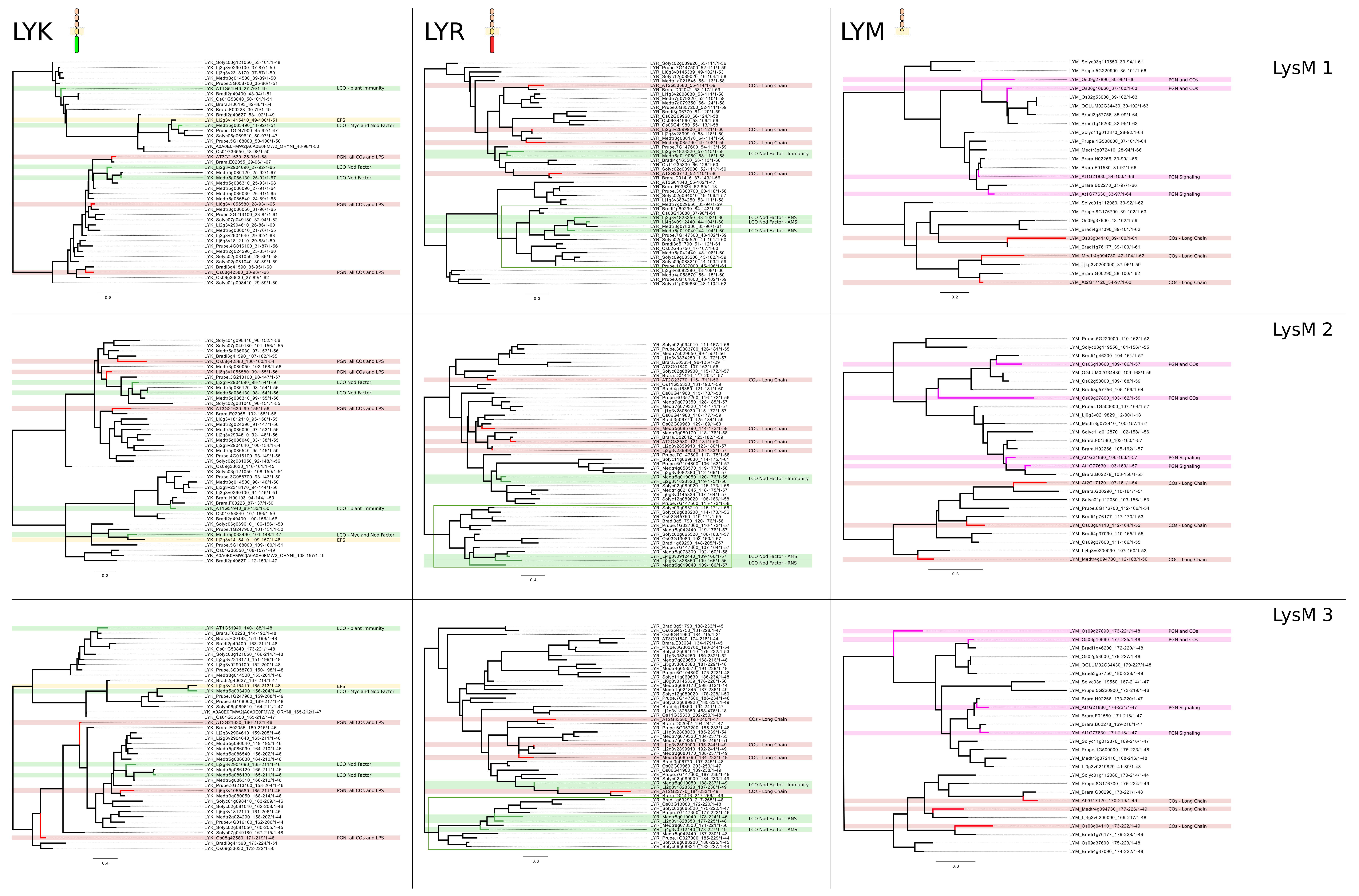
**Figure 3S. Phylogenetic trees of individual LysM domains from RLK/Ps classified by family.** The LysM1, LysM2 and Lysm3 domains of RLK/Ps were separated by family (LYK, LYR, LYM) and a maximum likelihood phylogenetic tree was constructed for each dataset. CO-IM: LysM triple bundles from Proteins involved in plant immunity triggered by chitin oligosaccharides. LCO-RNS: LysM triple bundles from proteins involved in rhizobium NF perception. PGN: LysM triple bundles from proteins involved in peptidoglycan sensing.

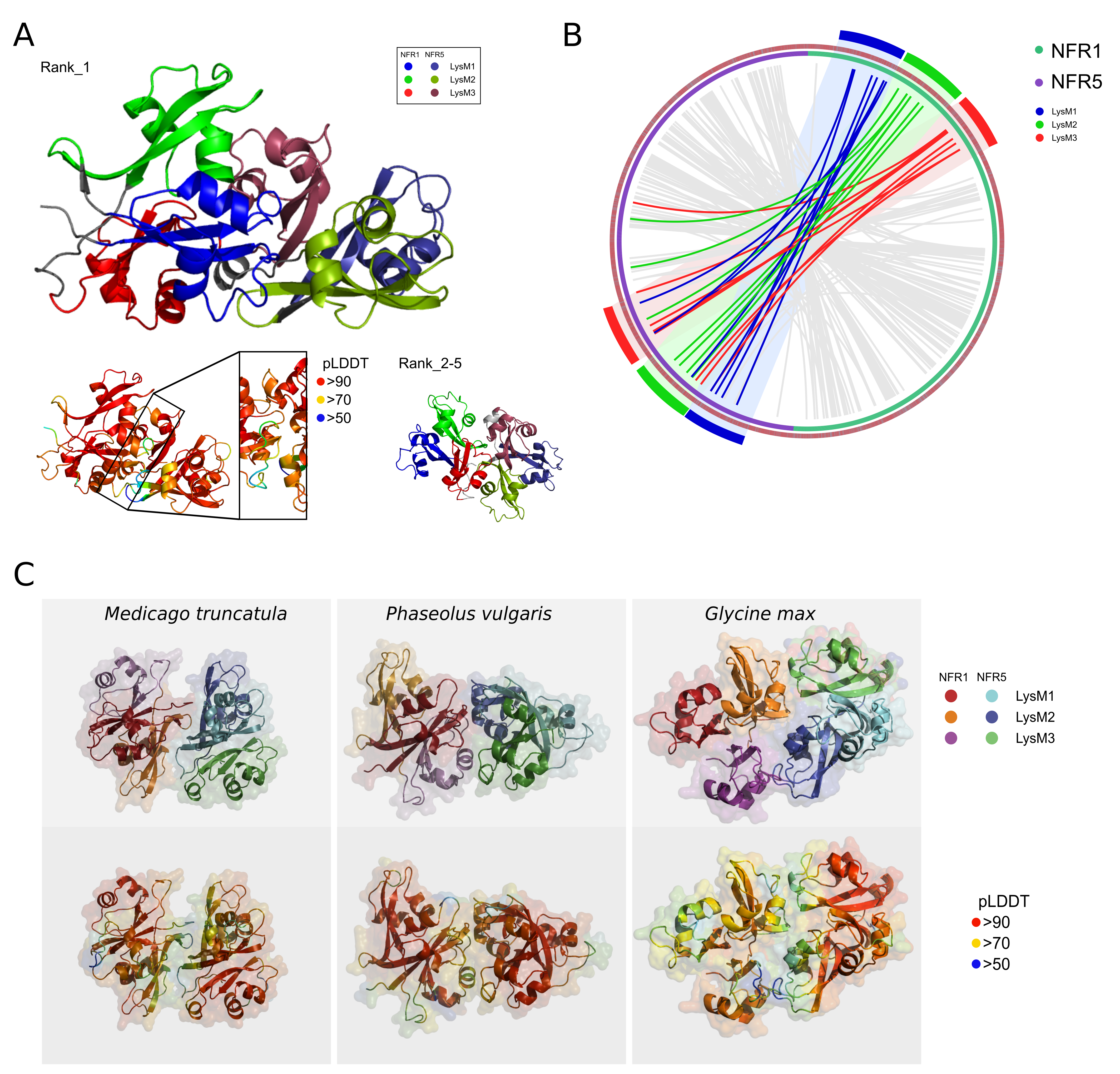
**Figure 4S. AlphaFold models of the interaction between NFR1 and NFR5.** ColabFold was used in multimer mode to predict the interaction between NFR1 and NFR5 homologs in legume plants. A. LjNFR1 and LjNFR5. B. I-COMS was used to detect coevolution signatures between LjNFR1 and LjNFR5. A circos plot is shown, with curved lines indicating coevolution between NFR1 and NFR5 positions. C. ColabFold models for BFR1 and NFR5 homologs from different model legume plants.

**Supplementary Tables**

**Table 1S. Sequences used in this study.** Description and accession numbers of all the sequences used.

**Table 2S. The results of using hmmsearch with the PFAM HMM to identify LysM domains in plant RLK/Ps from dataset used by Buendia et al. (2018).**

**Table 3S. The results of using hmmsearch with the ligand-specific HMM sets to classify plant RLK/Ps from dataset used by Buendia et al. (2018) by their ligand specificity.**
